# Practical TMS coils with maximum focality and various stimulation depths

**DOI:** 10.1101/2025.08.12.669952

**Authors:** Luis J. Gomez, Lari M. Koponen, Rena Hamdan, Yiru Li, David L. K. Murphy, Stefan M. Goetz, Angel V. Peterchev

## Abstract

Conventional transcranial magnetic stimulation (TMS) coils generate a diffuse and shallow electric field (E-field) in the brain. This results in limited spatial targeting precision (focality). Previously, we developed a methodology for designing theoretical TMS coils to achieve optimal trade-off between the depth and focality of the induced E-field, as well as the energy required by the coil. This paper presents methods for the practical design and implementation of such focal-deep TMS (fdTMS) coils. First, we consider how the coil’s shape affects energy requirements and design a curved “hat” former that enables a wide range of coil placements while improving energy efficiency compared to flat formers. Second, we introduce a hybrid layer winding implementation to improve energy efficiency by using multi-layer windings in some regions of the coil and a single layer in others. Using simulations with a spherical head model, we benchmark the focality of the fdTMS E-field in the brain and the scalp, as well as the required energy, against conventional TMS coils. We then implement two fdTMS coil designs using copper wire wound inside a 3d-printed plastic former. The E-field of the prototype fdTMS coils and two conventional figure-8 counterparts is measured with a robotic probe to validate the designs. The experimental measurements corroborate the simulations, demonstrating that the fdTMS coils produce a more compact E-field relative to standard figure-8 coils. One potential disadvantage of the fdTMS coil prototypes is wider spread of the scalp E-field and increased energy loss due to the additional windings. Nonetheless, the presented fdTMS coils could offer advantages for precise mapping studies, and the design framework could be leveraged for other coil optimizations.

## INTRODUCTION

Transcranial magnetic stimulation (TMS) is a non-invasive method for brain stimulation widely employed in neuroscience to investigate and probe brain function and connectivity. Furthermore, TMS has received FDA approval for treating of depression (1), as well as various other psychiatric and neurological disorders (2, 3). During a TMS session, a coil placed on the scalp and driven by brief strong current pulses induces an electric field (E-field) in the brain. Enhancing the focality and depth of this induced brain E-field provides greater flexibility and selectivity in targeting its effects. Consequently, numerous studies have attempted to design coils with improved focus and penetration depth (4-19). In our previous work, we developed a computational method for designing coils that achieve optimal trade-offs between focality, depth, and energy (fdTMS). Numerical studies demonstrated that coils designed using this framework can outperform the state-of-the-art figure-8 coils (20). In this paper, we further refine the design framework to consider practical coil implementation constraints, analyze depth–focality trade-offs compared to commercial coils, implement practical optimized coils with enhanced focality, and characterize them through electric field simulations and measurements.

Until recently, figure-8 type coils, consisting of two circular coils placed side-by-side, were considered the closest to achieving an optimal depth–focality trade-off (21). Computational coil design methods employing the stream function approach (22) have been used to design coils that significantly enhance energy efficiency (11, 19, 23, 24), reduce sound artifact (23), and produce more focused and deeply penetrating E-fields (11, 20). These studies have also indicated that to achieve a more focal stimulation, increased coil energy is required (11). Additionally, coil supports that are more conformal to the head can achieve better energy trade-offs than non-conformal ones (11, 23). However, perfectly conformal coils cannot be used across individual head shapes or different cortical targets, leading to the proposal of ‘hat’ shaped coils (23). In this paper, we present an optimized, practical hat-shaped coil support that leads to designs offering greater energy efficiency than flat supports. These hat-shaped coils are also ergonomically adaptable to various head sizes and shapes as well as scalp placements.

The half-maximum E-field threshold, which is defined relative to the maximum and thus is independent of it, has been commonly used in figures of merit to benchmark coils in terms of focality and depth of stimulation (21). Here we extend the study of focality by analyzing other thresholds and designing coils to optimize depth–focality trade-offs with respect to other E-field thresholds. We find that there are negligible differences between the coils designed with a half-maximum E-field threshold and other thresholds. These results indicate that the half-maximum region characteristics are effective as a figure of merit to rank coils in terms of the shape of the E-field distribution that is robust to variations in activation threshold.

The optimized fdTMS coils feature smaller windings concentrated near the center and form more intricate patterns compared to conventional figure-8 coils. For example, the typical fdTMS coil consists of a small figure-8-like coil with four additional ‘cancellation’ loops and two large ‘biasing’ loops on the sides. The relatively small sizes of the figure-8-type loops and the wire’s cross-sectional area requirements for energy efficiency necessitate the use of multiple layers to achieve inductances compatible with coil drivers. These additional layers, in turn, increase the coil energy requirements. We developed a synthesis technique for the coil windings to include multiple layers in critical regions where the inclusion of required additional turns is not feasible, and single layers elsewhere. This approach mitigates the increase in energy requirements resulting from the use of multiple layers. Finally, we introduce a semi-automatic procedure for generating a 3d coil former that has grooves where the coil windings reside, with added features for safety and comfort of the subject. The coil E-fields are simulated and measured in a spherical phantom and the fdTMS coils’ performance is benchmarked relative to commercial coils.

## METHODS AND MATERIALS

In this section, we describe our proposed approach for designing and fabricating fdTMS coils. Firstly, we introduce the coil design parameters and figures of merit used to assess coil performance. Detailed methodologies for optimizing these parameters were previously presented (20). Secondly, we describe the methods used to modify the optimal design, making it compatible with our coil fabrication procedures. In the third part, we outline the procedures for coil fabrication and provide details about the validation measurements of the coil’s E-field. Lastly, we provide additional details about figures of merit used for computational comparisons with conventional coils.

### Coil design parameters

The coil windings are assumed to reside on a surface Ω that is approximated by a triangle mesh consisting of *n*_*p*_ nodes. The winding paths are chosen such that when driven by a TMS coil driver they will approximate the E-field generated by a surface current on Ω (this process is given in the next section). To determine the optimal current, it is assumed that it is in the span of the *n*_*p*_ seed currents 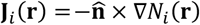, where *i* = 1,2, …, *n*_*p*_, **r** = (*x, y, z*) denotes Cartesian position, and *N*_*i*_ (**r**) are nodal finite elements on the triangle mesh (22). In other words, surface current distributions **I**(**r**, *t*) are defined on a coil surface Ω (Figure 1) as

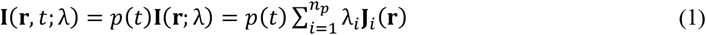

where 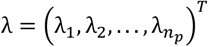 is a vector of weights, each λ_*i*_ (where *i* = 1,2, …, *n*_*p*_) is a real number, and *p*(*t*) = sin (ω*t*) and ω = 3000 · 2*π*. Note that *p*(*t*) was assumed to be time-harmonic to simplify the exposition. However, because of the relatively low-frequency content of TMS pulses, the results apply to other current waveforms as well.

**Figure 1.**
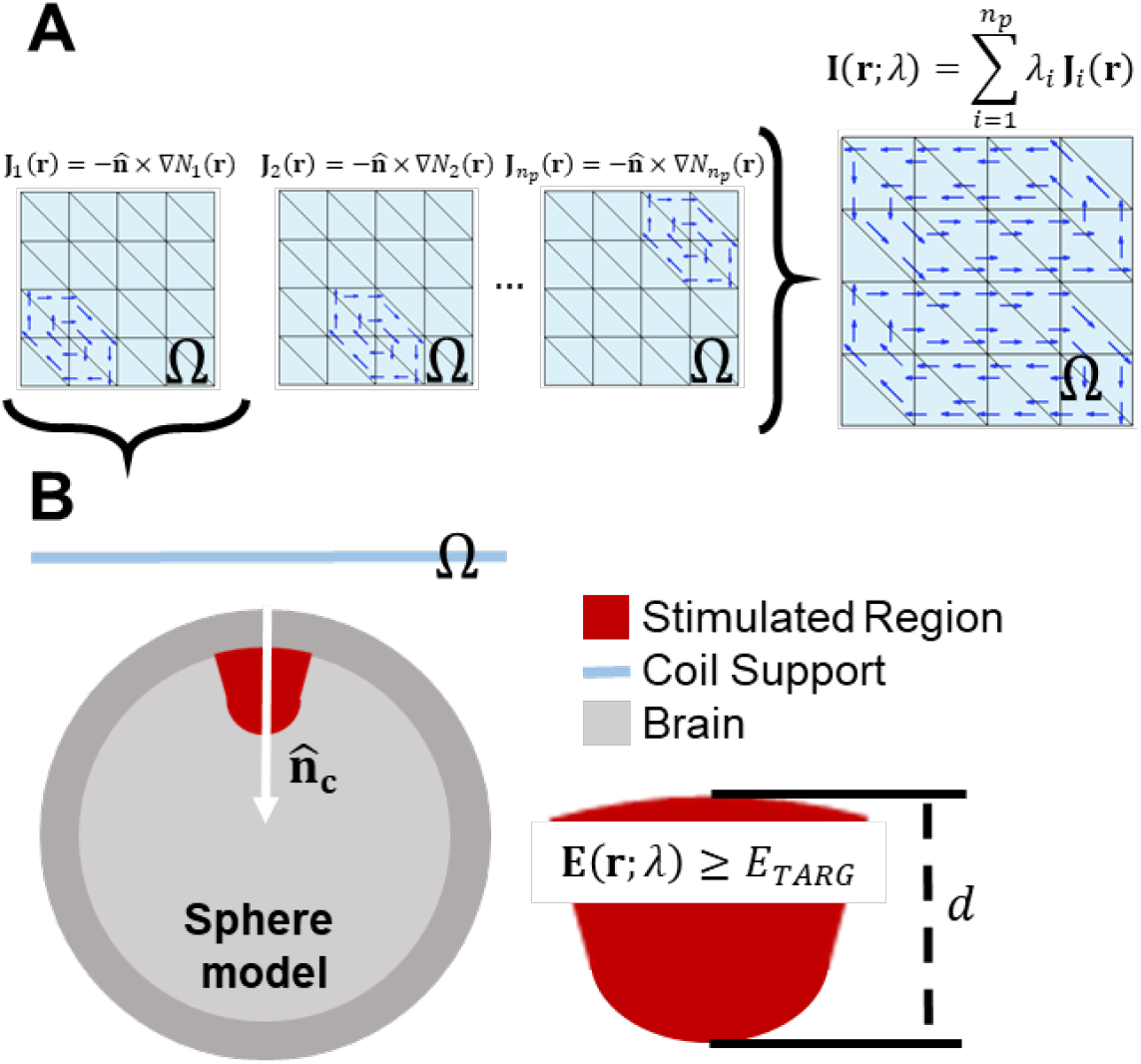
Coil current density parameters and E-field figures of merit definition. (A) Each seed current corresponding to internal nodes of the coil support triangle mesh is linearly combined to generate surface current distributions. (b) The E-field generated in a spherical head model by each coil is computed, and stimulation depth and volume figures of merit are extracted.

The above seed currents are known to span all piecewise linear currents with zero divergence on the triangle mesh (25), thereby forming an adequate basis for approximately including all admissible E-fields generated by non-dissipative surface current distributions on Ω.

### Coil performance figures of merit

The fdTMS coil optimization takes as input a triangle mesh (or parametrization) of the surface Ω and a set of seed current distributions **J**_*i*_ (**r**), where *i* = 1,2, …, *n*_*p*_. Then, it finds Pareto optimal currents **Ir**, *t*; λ_*opt*_ that achieve optimal trade-offs with respect to stimulation energy, depth, and volume. The coil figures of merit are defined as follows:

#### i) Minimum stimulation volume

The stimulated volume *V* is defined as

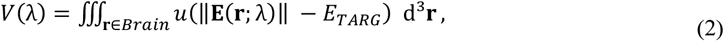

where **E**(**r**; λ) denotes peak E-field at location **r** induced by the surface current **I**(**r**, *t*; λ), ‖·‖ denotes vector magnitude, *u*(*x*) is a unit step function, the integration is over the brain region (denoted *Brain*), and *E*_*TARG*_ is the E-field at the targeted depth location. The value of *u*(‖**E**(**r**; λ)‖ − *E*_*TARG*_) is one if ‖**E**(**r**; λ)‖ is above the stimulation threshold and zero otherwise. As a result, Eq. (2) measures the volume of the region above threshold.

#### ii) Maximum depth of stimulation

The depth of stimulation *d* is defined along a line **s**(*l*) chosen as a line that intersects at and is perpendicular to the center of the surface current support, i.e. 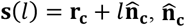 is the head unit normal pointing inwards at the point nearest to the coil center. In accordance, the stimulation depth is

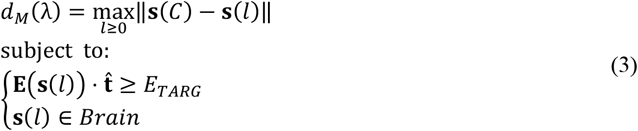

where **s**(*C*) denotes the point on the cortex closest to **r**_**c**_. For example, Figure 1B depicts the coil placed centered about and oriented perpendicular to the z-axis. In this case, the line in blue denotes 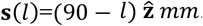. Furthermore, markers are included at 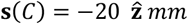 and the lowest point with E-field above threshold **s**(*C* + *d*_*M*_ (λ)). The value of *d*_*M*_ (λ) is the distance between these two points and the choice of 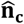 pointing toward the brain results in **s**(*C* + *d*_*M*_ (λ)) being the deepest point stimulated.

#### iii) Minimum energy

TMS pulses have relatively low-frequency temporal variation and their induced magnetic field is negligibly affected by the presence of the head. The magnetic energy stored in the current distribution can be computed using the Biot-Savart law (26) as

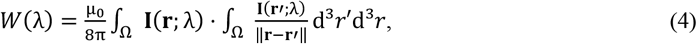

where μ_0_ is the permeability of free space.

In addition to the aforementioned figures of merit, we combine the *V* and *d* to define Spread (*S*) as the average transverse surface area of the stimulated region, calculated as *S* = *V*/*d*_*M*_ (21, 27). Reducing either *V* or *S* is equivalent to an increase in focality. Moreover, safety considerations impose limits on the maximum E-field strength that brain tissue can withstand. For a given α, we assume that E-field strength exceeding α*E*_*TARG*_ in the brain is unacceptable. Therefore, currents in the span of the modes that result in an E-field that exceeds α*E*_*TARG*_ in the brain are excluded from the admissible designs. In prior research, a common selection for *α* is *α* = 2. In this scenario, *V, d*_*M*_, and *S* are equal to the figures of merit *V*_½_, *d*_½_, and *S*_½_ = *V*_½_/*d*_½_, respectively (20, 21, 27, 28). Consequently, this leads to *V* representing the sub-volume of the brain where the E-field equals or exceeds half of its peak value, and *d*_*M*_ representing the greatest depth where the E-field equals or exceeds ½ of its peak value.

In many instances the peak E-field on the cortex is less than twice *E*_*TARG*_. Furthermore, TMS pulse current can be increased or decreased thereby allowing for the coil to stimulate deeper or shallower regions. As such, the energy and spread of a fixed coil is not a constant but a function of targeted depth. To account for this in the coil benchmarks we additionally considered the stimulation volume, spread, and energy for each coil as a function of targeted depth, i.e. *V*_*d*_ (*d*), *S*_*d*_ (*d*), and *W*_*d*_ (*d*), respectively, where *d* ∈ (0, *d*_*M*_]. *V*_*d*_ (*d*) and *S*_*d*_ (*d*), and *W*_*d*_ (*d*) are computed by driving the coil with a current that results in an E-field of *E*_*TARG*_ at *d*. Furthermore, targeted depths that result in a peak cortical E-field above α*E*_*TARG*_ are considered unreachable and excluded from the evaluation of *V*_*d*_ (*d*), *S*_*d*_ (*d*), and *W*_*d*_ (*d*).

Finally, we consider the design of coils with the alternative choice of 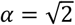 used in some publications (11). We find that *V*_*d*_(*d*), *S*_*d*_(*d*), and *W*_*d*_(*d*) for designs with either *α* = 2 or 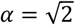 achieve similar trade-offs for shallow depths, and the ones designed with *α* = 2 achieve superior performance for larger depths. As such, we use and recommend *α* = 2 to design fdTMS coils.

### Coil winding generation

Here we discuss how we convert the current density 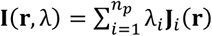 into coil windings. Our approach starts by adopting the standard coil winding generation approach that defines the stream function of the current 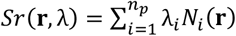, where 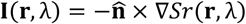. The contour lines of the stream function point in the direction of the surface current and trace out regions of constant current. Furthermore, since the current is dependent on the gradient of the stream function, an approximation of the surface current is made by placing wires on contour lines that are an equal elevation distance apart (i.e. equispaced contour intervals). For example, if we choose *M* equispaced contour intervals, then, the wires are placed at contour elevation levels of

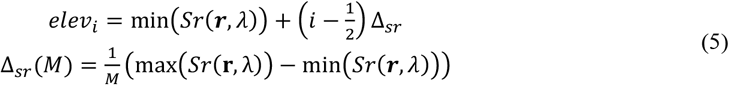

where *i* = 1, …, *M*. These wires are then typically connected serially to generate a coil design. This approach has been shown to yield coil designs that match the surface current distribution and by proxy E-fields. The value of *M* is typically chosen to result in a coil design that matches a desired inductance.

For fdTMS coil designs the contour lines typically concentrate near the coil center and it is impossible to attain the TMS driver required inductance (≥ 8.5 μH) without using multiple layers of wire. Adding multiple winding layers results in a design that is less energy efficient than a single layer design. To achieve the desired inductance while accommodating enough contour intervals to achieve a desired inductance we adopted two strategies.

One evaluated strategy was to add constraints to the optimization to ensure that the stream function would allow enough concentric loops to fit the required number of turns of the windings. This is done by introducing a maximum constraint in the L-infinity norm of the solution as this has been shown to yield designs with more spread-out concentric windings (24). For a given choice of *M*, the closest any two windings can theoretically be is *dist*_*min*_ = Δ_*sr*_ (*M*) max ‖**I**(**r**, λ)‖. We set the additional constraints to our fdTMS design framework to guarantee that *dist*_*min*_ ≥ *W*_*wire*_, where *W*_*wire*_ is the wire width.

We iteratively implemented this constraint by first running the fdTMS framework and adding constraints where the condition was violated. The value of Δ_*sr*_ (*M*) is determined using the stream function of the previous design iteration. The constraint of max ‖**I**(**r**, λ)‖ is approximated by 16 linear constraints by using the same approach that was used for the E-field constraint in (11). This procedure converged after 2–5 iterations for all cases tested here. For any given *M, didt*_*min*_ can only be reduced marginally while maintaining the same performance. As a result, these additional constraints still required the use of multiple layers.

The second evaluated strategy was to adopt a hybrid approach where the coil has multiple layers only on critical regions where the concentric windings are densest. Specifically, the majority of fdTMS coils consist of three distinct types of sub-coils: figure-8, biasing, and cancellation windings. The biasing and cancellation windings allow enough space to be implemented using one layer, while the figure-8 winding requires three layers. We first partition the coil surface Ω into the three subregions containing the figure-8, biasing, and cancellation loops, respectively. For each part we specify the number of concentric loops in our implementation and number of layers. Each concentric loop consists of wire placed at a contour line. As a result, the design is specified by the contour line elevation levels and layers for each coil sub-region. The contour line height levels are determined by running an optimization to minimize the error between E-fields generated by the coil and the optimal surface current.

To numerically achieve the above objective, the E-fields of the coil and the optimal surface current are sampled uniformly at 46,532 points on a spherical shell and assembled into 46,532 × 3 matrices **E**_**Coil**_ and **E**_**Surf**_, respectively. Furthermore, to regularize the optimization we add an energy penalty equal to 10^−5^‖**E**_**ideal**_‖_*F*_ W_coil_, where ‖**E**_**ideal**_‖_*F*_ is the Frobenius norm of **E**_**ideal**_ and W_coil_ is the energy required by the coil to generate an E-field that optimally matches **E**_**ideal**_. Finally, a constraint is added to ensure that only designs with concentric loops 2.2 mm apart are admissible. The final optimization is done using MATLAB’s fmincon to minimize the function ‖**E**_**Coil**_ − **E**_**ideal**_‖_*F*_ + 10^−5^‖**E**_**ideal**_‖_*F*_ W_coil_.

The approaches described above result in designs consisting of disconnected loops. The loops are manually connected serially. The connections were chosen with the following considerations in mind: branching from concentric windings should be done far from the coil center, branching between coils is both chosen where neighboring coils are closest, and the curvature of the transitions between coils should be small enough to enable practical winding of the coils.

To benchmark the procedure described above, we generate winding patterns for fdTMS coils that reach or exceed the penetration depths of existing TMS coils. Specifically, the above procedure is applied to design fdTMS coils with *d*_½_ ≥ 1.01, 1.31, and 1.57 cm, which are the *d*_½_ values of MagVenture B35, B65 and B80 coils, respectively. In each case, we choose the number of turns to result in an inductance ≥ 8.8 *μH*.

### Coil shape and former design

The hat-shaped coil support is derived from five MRI-based head models (29) using SimNIBS (30). Specifically, we densely sample coil placements on the motor strip of each subject. Then, the most conformal radially symmetric shape that would allow orienting and placing the coil on all the candidate motor strip positions is chosen. The general shape is given in Figure 2B. To compare the coil shape’s effect on performance, we additionally considered sphere (Figure 2A), half-sphere (Figure 2C), and square (Figure 2D) coil supports.

**Figure 2.**
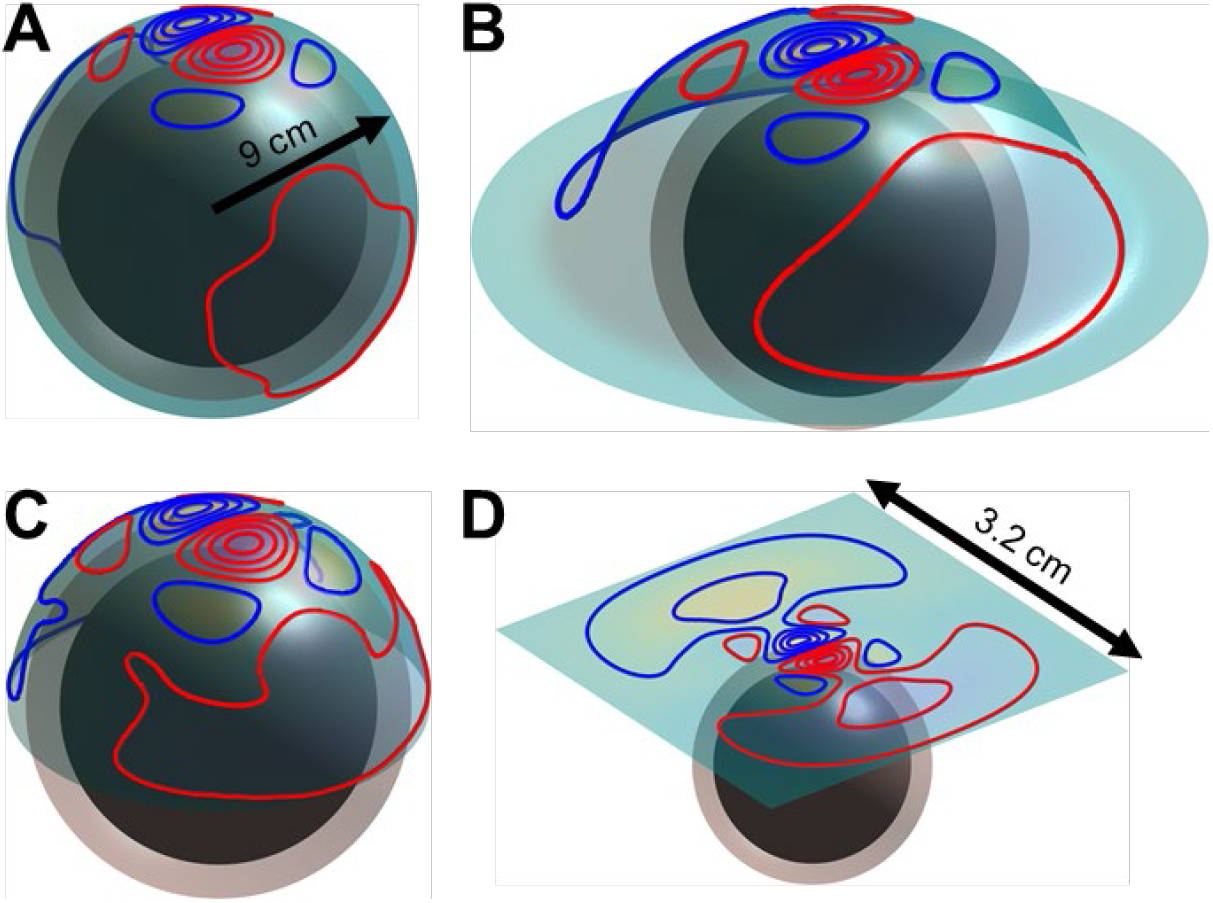
Evaluated fdTMS coil supports (surface shapes): (A) sphere, (B) hat, (C) half-sphere and (D) square support placed over the spherical head model. The inner-most sphere is the brain region.

We generate several meshes by extruding the coil about its normal direction and converting the filamentary wire representations to thick wire meshes (Figure 3). First, a thin version of the coil support and thick and tall winding meshes are merged (Figure 3A). This first step allows us to have tall enough coil channel grooves without requiring a heavy coil support. Second, ribbed windings are subtracted from the merged mesh (Figure 3B). The resulting mesh has grooves where the windings will reside. The narrower segments of the grooves mechanically hold the wire in place, whereas the wider stretches of the grooves reduce friction during the wire insertion and provide space to inject epoxy to bond the wire to the former.

**Figure 3.**
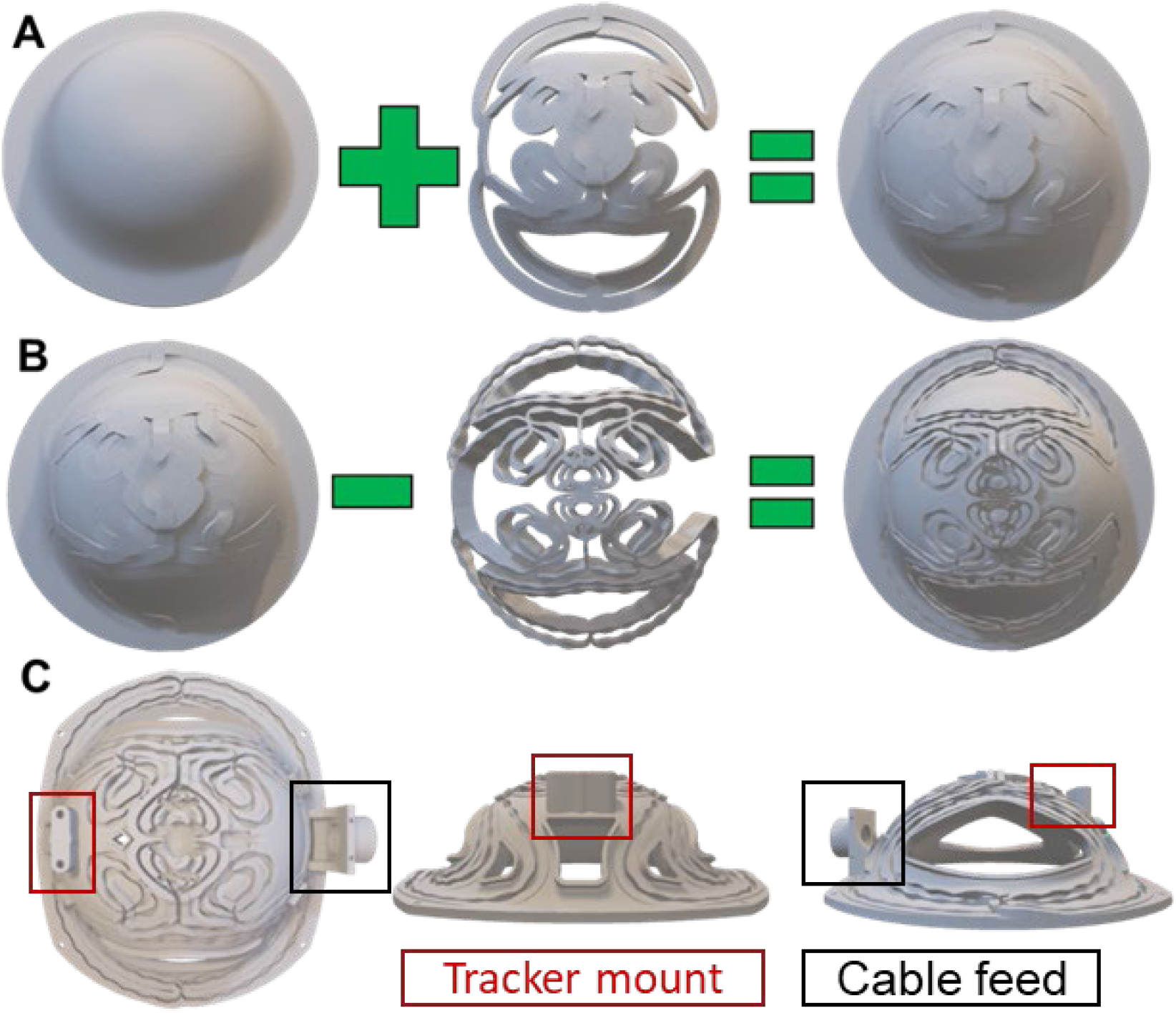
Fabrication steps of fdTMS coil hat-shaped former. (A) Coil support mesh is merged with a thick and tall wire mesh. (B) Ribbed wire mesh is subtracted from the merged mesh in (A). (C) Three views of the final mesh with additional mounting supports for cable feed and neuronavigation tracker, and holes cut out for ventilation and head visibility.

### Coil fabrication

We built two fdTMS coils—one to target a depth of 1.31 and the other to target a depth of 1.57 cm to match the depth characteristics of the Cool-B65 and B80 coil, respectively (MagVenture A/S). The coil former (Figure 3C) is 3D printed using selective laser sintering of Nylon PA 12. The top and bottom surfaces of the former are then sprayed with electrical sealant to provide additional insulation. A 12 AWG (2.05 mm diameter) round magnet wire is wound in the former grooves following the winding path. During the winding process, small drops of hot glue are used intermittently throughout the path to secure the wire in place. Once the entire coil is wound, quick-set epoxy is injected into the grooves and allowed to fully cure for 24 hours. Finally, the two magnet wires exiting the coil are soldered to copper connectors that are then attached to a cable assembly compatible with MagPro TMS devices (MagVenture A/S). To monitor for safe coil operation, a temperature sensor wired to the cable assembly is affixed to the former near the center of the coil winding. The coil windings and cable conductors are tested at 10.7 kV relative to the external coil surface to ensure safe electrical insulation per standard IEC 60601-1 (31)

### Coil electric field measurement

To validate the design of the fdTMS coils, their E-field as well as the E-field of conventional figure-8 coils (MagVenture Cool-B65 and B80) are measured across a hemispherical shell using triangle probes mounted on a robotic stand (32). The voltage induced in two aligned triangle probes with heights of 7 cm and 6 cm is measured, corresponding respectively to the E-field strength on the surface of a conductive sphere with 7 cm radius, matching the brain compartment of the spherical head model used in the fdTMS coil design and evaluation, and the E-field at 1 cm depth from the brain surface. A custom 3D printed stand serves to center the TMS coil directly above the probe center of rotation and to hold the coil and probe setup in place. The TMS coil surface is positioned 8.7 cm from the center of the probe, corresponding to 1.7 cm from the brain surface. TMS pulses are delivered to the coil using a MagPro R30 device (MagVenture A/S) at 27% of maximum stimulator output. The stimulator is configured to send a sequence of biphasic pulses at a rate of 0.5 pulses per second in 10 trains of 364 pulses each with an intertrain interval of 1 second. The probe is controlled using Arduino and a MATLAB code that is set to record E-field data along a specified path. A total of 3,620 data points is recorded and used to generate surface E-field maps.

### Head model

A common head model consisting of a homogenous sphere with radius of 8.5 cm is used for the fdTMS coil design and evaluation (11, 21). The head model consists of two concentric spheres each centered about the origin and having radii of 7.0 cm and 8.5 cm, respectively. The inner sphere corresponds to the brain, and the outer shell—to the CSF, skull, and skin. Since TMS does not induce any radial currents in a spherical conductor and the E-field is independent of the specific conductivity according to the quasi-static modeling approximation, a single conductivity of 0.33 S/m is used for the whole sphere (33). For each simulation (Figure 2A–D), the center of the coil in Cartesian coordinates is **r**_**c**_ = (0,0,0.09) and 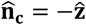. Correspondingly, depth is measured along the −*z* direction starting from *z* = 7.0 cm. Analytical expressions for the E-field generated inside the spherical head model are given in (11) and used to determine the E-field generated by the surface currents in the context of the fdTMS coil design optimization.

### Coil performance evaluation with finite element simulation

Using the same definitions for the spherical head model, stimulation volume *V*, energy *W*, and depth of stimulation *d*, we characterize the stimulation volume and energy of fdTMS and, for comparison, standard figure-8 coil models over a range of target depths using finite element simulations in SimNIBS (30). The sphere head model is recreated in SimNIBS with 30.7 million tetrahedra. The fdTMS coil definitions are imported into SimNIBS by generating a voxel grid of primary E-field samples and storing them as NIfTI files, and the native SimNIBS models for three conventional TMS coils are used. The E-field strengths on the outer (scalp) and inner (cortical) surfaces of the spherical model are calculated at the center of each triangle that constitutes the corresponding surface mesh during finite element analysis. We then calculate the peak E-field strength on the cortical surface over a range of depths and characterize the distribution of the E-field on the model scalp surface.

## RESULTS

### Energy vs. focality for various coil supports

Here we analyze trade-offs between focality and required energy for various target depths and coil topologies. Specifically, we compare the energy vs. focality of the hat coil relative to the sphere, hemisphere, and flat coils of our previous work (20) and of our current work including wire thickness. Figure 4 shows energy versus spread curves for target depths *d*_½_ ≥ {1.01, 1.31, 1.57} cm. Coils with spherical support are the most conformal and they are more focal than the others for matched depth and energy. The coils with hat support exhibit performance that is better than the square support coils and slightly worse than the hemispherical coils. The efficacy of conformal relative to non-conformal coils increases with depth: The flat coils preform nearly as well as the hat shaped ones for *d*_½_ ≥ 1.01 cm (Figure 4A,D), whereas flat coils are significantly inferior for *d*_½_ ≥ 1.57 cm (Figure 4C,F). Independent of depth, the fdTMS coils outperform the conventional coils by either energy for matched spread or spread for matched energy, consistent with our prior findings (20).

**Figure 4.**
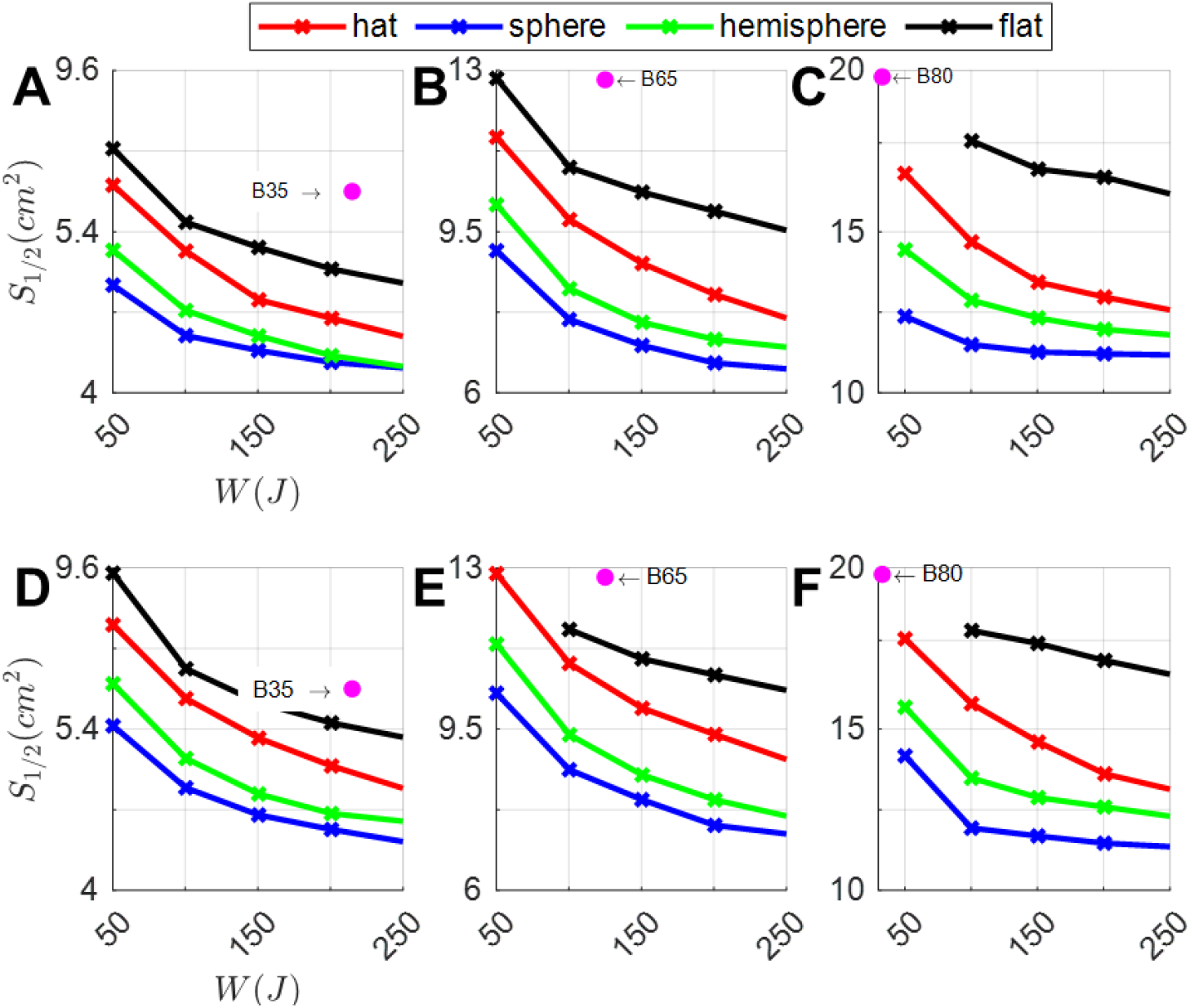
Trade-off between E-field spread and coil energy for fdTMS coils with different support surfaces (see legend) and (A)–(C) winding height of 6 mm or (D)–(F) filamentary winding height. The columns correspond to depth of (A),(D) *d*_½_ ≥ 1.01 cm; (B),(E) *d*_½_ ≥ 1.31 cm; and (C),(F) *d*_½_ ≥ 1.57 cm, matching commercial coils MagVenture B35, B65, and B80, respectively, whose performance is denoted with pink dots for comparison.

We additionally considered designs assuming that 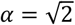. Figure 5 shows energy versus 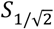 curves for target depths 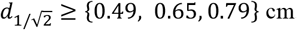. Same as before, spherical support (i.e. more conformal) and hat coils outperform square flat fdTMS coils. Figures 4A–C and 5A–C display results for wires having 6 mm height and Figures 4D–F and 5D–F—for filamentary wires. The filamentary wires always outperform their corresponding non-zero height wire results, indicating that height of the coil should be minimized to ensure best spread and energy performance, to the extent possible within the constraints of conductive power loss and associated coil heating.

**Figure 5.**
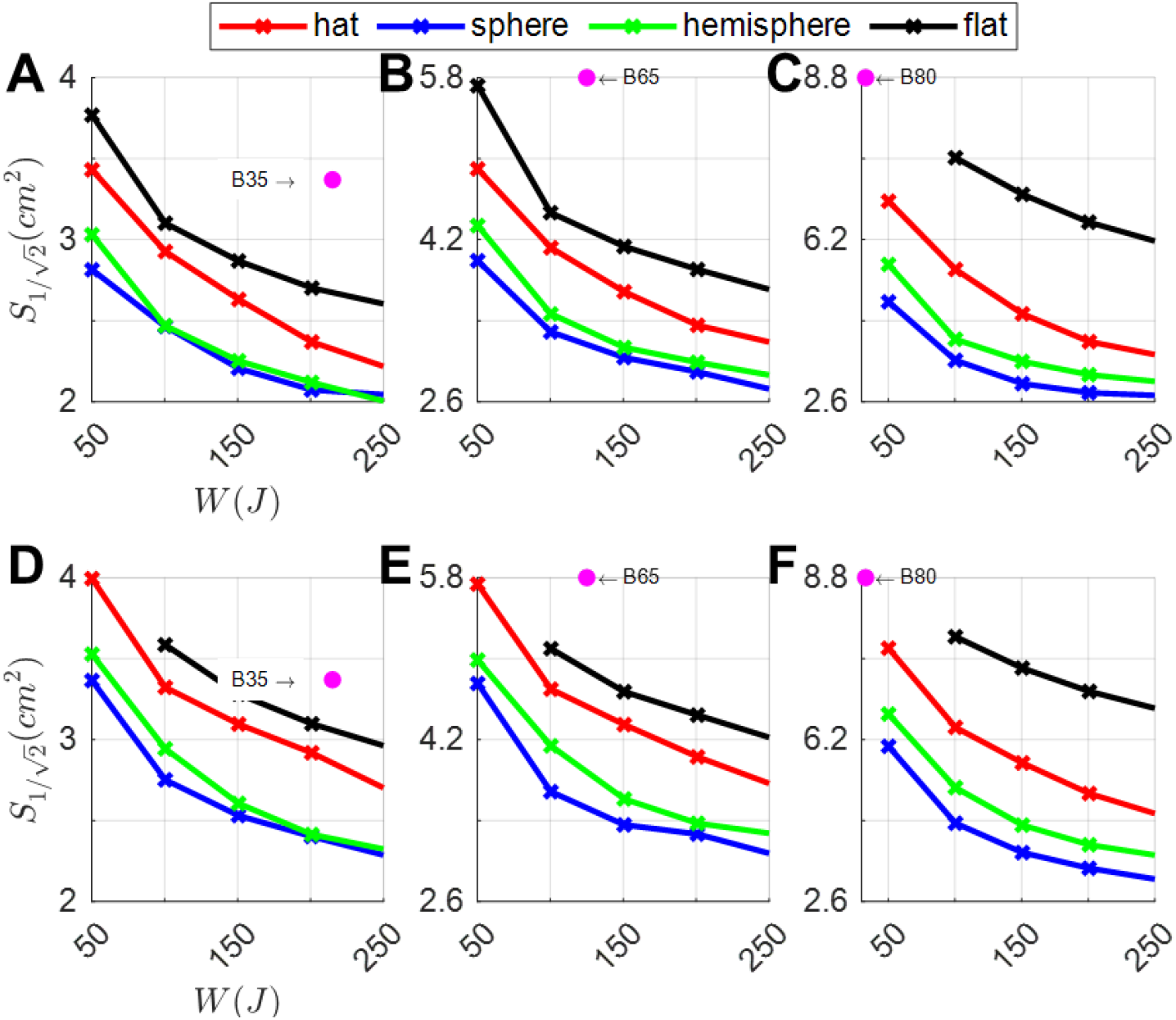
Trade-off between E-field spread and coil energy for fdTMS coils with different support surfaces (see legend) and (A)–(C) winding height of 6 mm or (D)–(F) filamentary winding height. The columns correspond to depth of (A),(D) 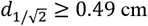; (B),(E) 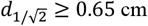; and (C),(F) 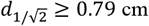, matching commercial coils MagVenture B35, B65, and B80, respectively, whose performance is denoted with pink dots for comparison.

### Spread as a function of depth

The amplitude of the coil driving current can be changed to generate a peak cortical E-field that is an arbitrary percentage of the stimulation threshold. For coil design, however, the peak cortical E-field is constrained to α times the E-field strength at the target, *E*_*TARG*_. Here we consider the depth–focality trade-off as a function of stimulation depth or, equivalently, as we allow α to vary from 0 to 2. Figure 6 shows that designs with 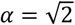 (i.e. peak cortical E-field of 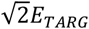) and *α* = 2 (i.e. peak cortical E-field of 2*ETARG*) have similar performance for shallow depths. In contrast, the fdTMS coils with *α* = 2 significantly outperform 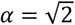 designs for deeper targets. Furthermore, as the energy of the coils increases, these performance differences become more significant. This is because the fdTMS coils tend to have large regions of their E-field with strength slightly below threshold. As such, when the fdTMS coils are operated with a peak E-field stronger than *αETH*, the stimulated region will be diffuse, reducing focality. Correspondingly, the choice of *α* = 2 is more robust than 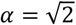 even if the coil will be operated to generate a peak cortical E-field less than twice the field strength at the target.

**Figure 6.**
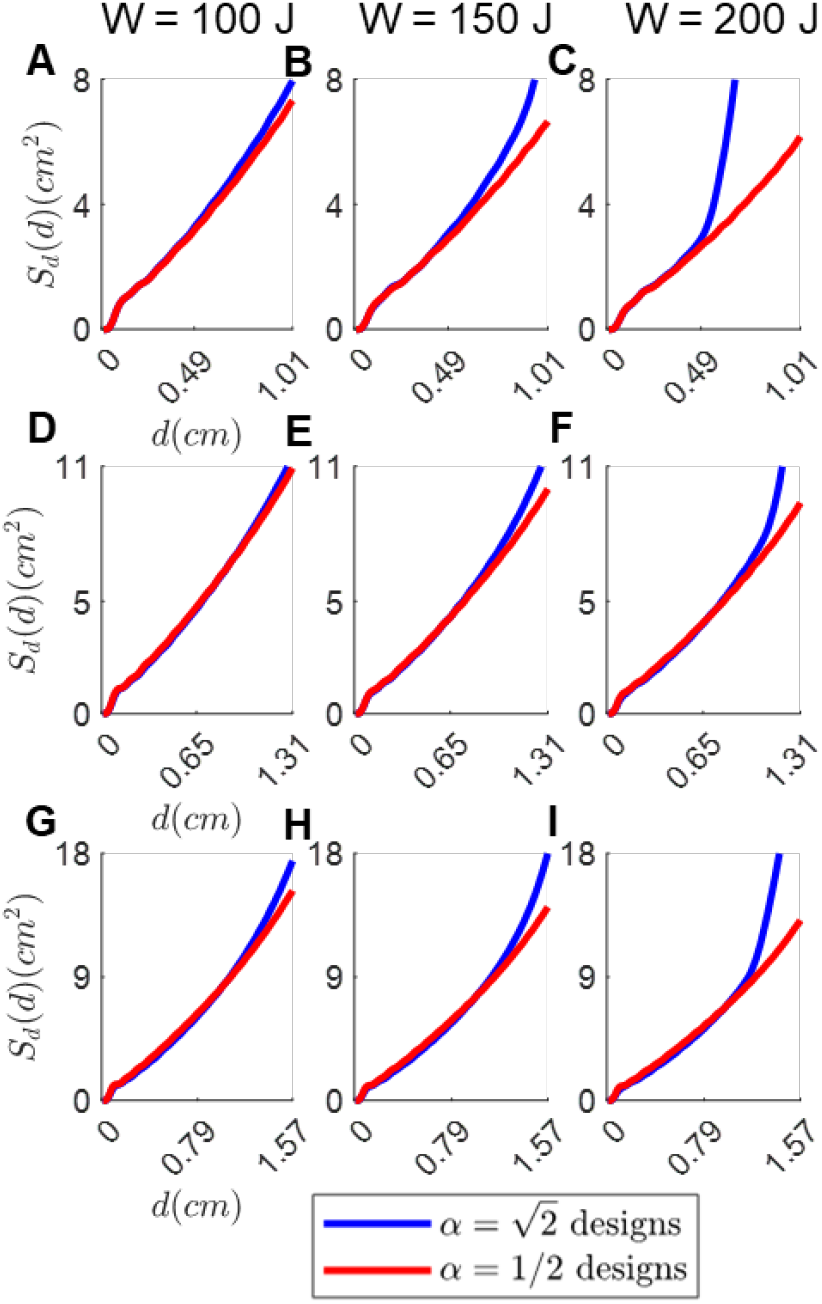
The E-field spread as a function of dept for coils designed with *α* = 2 and 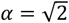. Results are arranged column-wise in increasing energy: (A), (D), and (G) W = 100 J; (B), (E), and (H) W = 150 J; and (G), (H), and (I) W = 200 J. Rows compare fdTMS coils with fixed depth of stimulation (A)–(C) 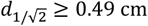 and 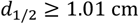, (D)–(F) 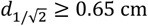 and 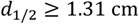, and (G)–(I) 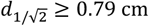 and 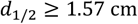.

### Choosing number of concentric windings

For practical coil drive currents and efficient energy transfer, TMS coils have inductance L ≥ 8.5 μH (34). Furthermore, to prevent heating and minimize the resistance of the coil, the wire cross-section is ≥ 8 mm^2^. The fdTMS coils typically have winding loops that concentrate in the coil center; as such, multiple layers must be used to achieve a high enough inductance.

We considered two implementations of the fdTMS coils: one using two layers and one using hybrid layers (Figure 7). The two-layer implementation has 2 mm wide, 4 mm tall wire, and 5 concentric loops (i.e. *M* = 10) when discretizing the stream function. This was the maximum number of concentric loops that could accommodate 2 mm wide wire. The hybrid-layer approach used three layers to implement the “figure-8” winding in the coil center and one layer for all other windings, with 3.0 mm by 3.0 mm wire cross-section. Table 1 summarizes key parameters of the hybrid-layer and two-layer coil designs. The hybrid coils achieved focality that is slightly inferior to the two-layer designs. However, the hybrid-layer designs used less energy than their two-layer counterparts (the energy limit target was 200 J); therefore, we chose to implement the hybrid-layer coils.

**Table 1.** Performance figures of merit for two-layer and hybrid-layer fdTMS coil implementation approaches.

|  | Two-layer |  |  | Hybrid-layer |  |  |
| --- | --- | --- | --- | --- | --- | --- |
| $d_{1/2}$ | $\geq 1.01$ cm | $\geq 1.31$ cm | $\geq 1.57$ cm | $\geq 1.01$ cm | $\geq 1.31$ cm | $\geq 1.57$ cm |
| $S_{1/2}$ | 5.9 cm<sup>2</sup> | 9.23 cm<sup>2</sup> | 13.2 cm<sup>2</sup> | 6.3 cm<sup>2</sup> | 9.7 cm<sup>2</sup> | 14.4 cm<sup>2</sup> |
| $W$ | 215 J | 197 J | 197 J | 168 J | 163 J | 169 J |
| $L$ | 8.1 μH | 11.9 μH | 20.5 μH | 8.8 μH | 8.8 μH | 13.5 μH |

**Figure 7.**
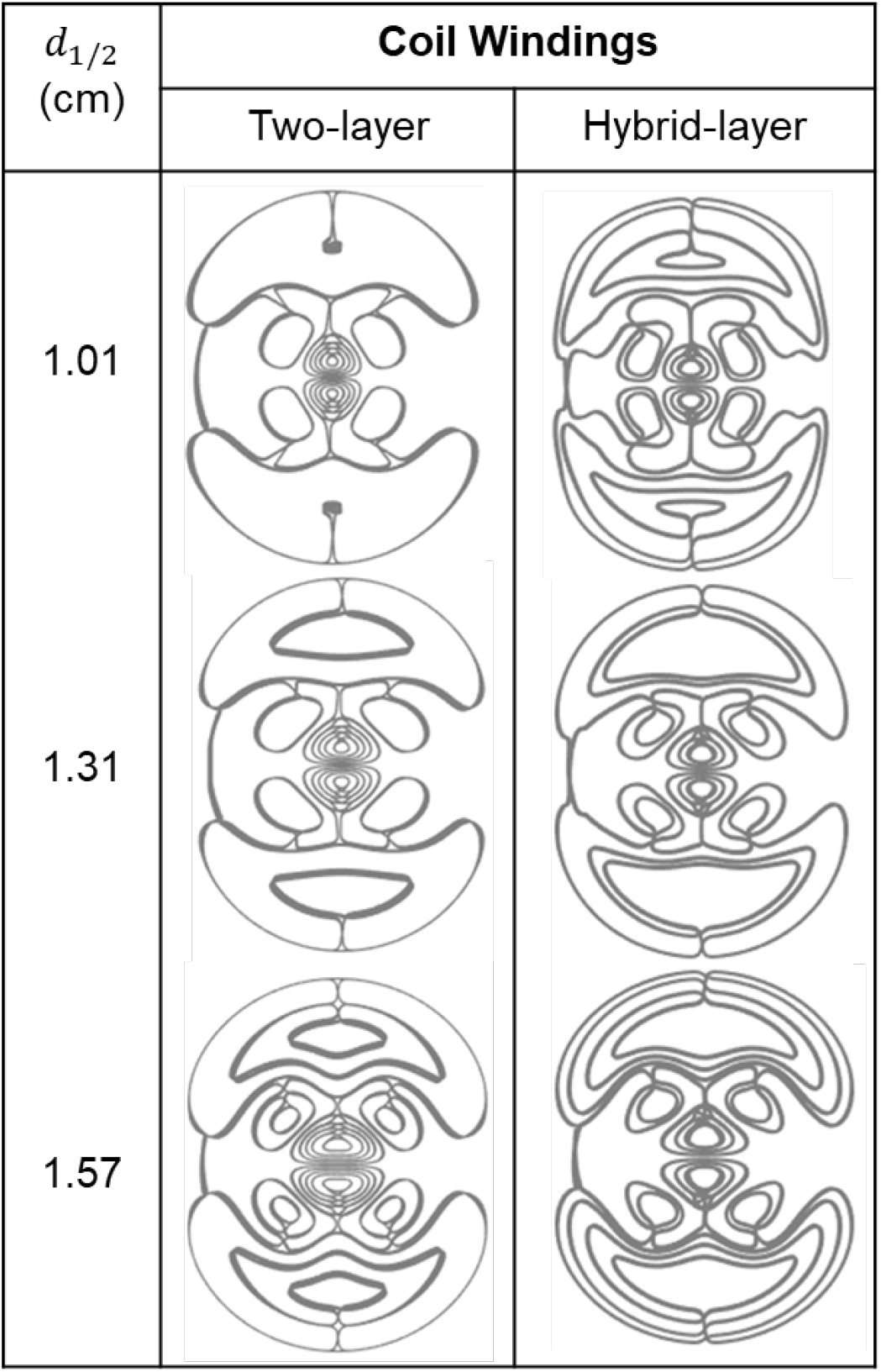
fdTMS coil designs with two-layer and hybrid-layer (three layers in the center, one in the periphery) windings.

### Electric field characterization

Figures 8 and 9 show the E-field distribution for conventional figure-8 coils and fdTMS coils. For the fdTMS coils, the region with E-field exceeding that of the target is more compact than that of the figure-8 coils for matched depth. Additionally, in Figure 9 the region above threshold for the fdTMS coil as it penetrates the head appears more compact and cylindrical than that of the figure-8 coils. These results indicate that lower *S*_½_ of the fdTMS coils does result in a more compact E-field distribution.

**Figure 8.**
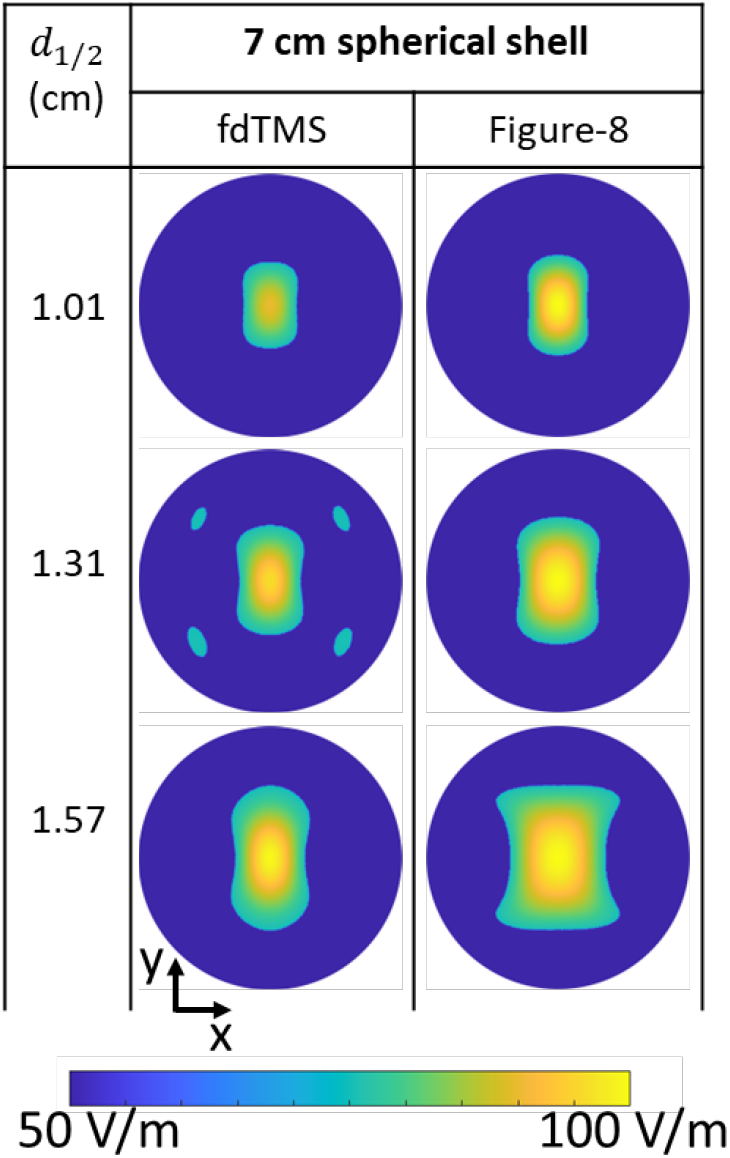
Coil E-field distribution on a 7 cm hemispherical shell for fdTMS and conventional coils.

**Figure 9.**
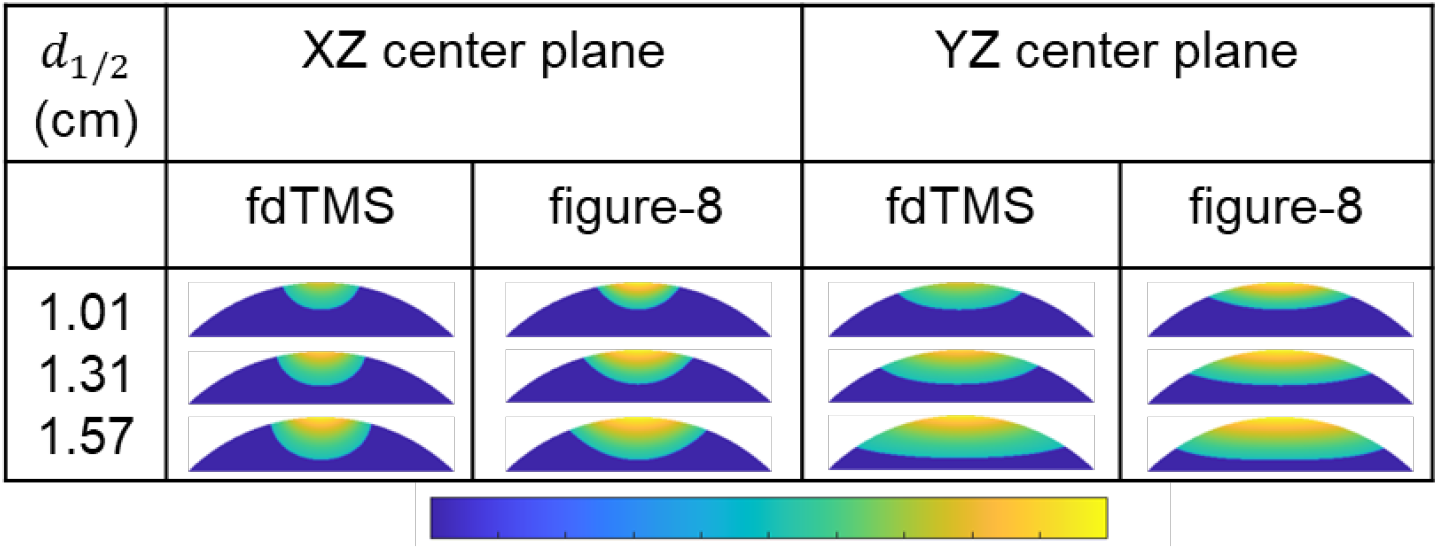
Coil E-field distribution on center cut planes for fdTMS and conventional coils.

Figure 10 shows the suprathreshold stimulation (*E* > *E*_*TARG*_) volume and peak E-field in the brain versus depth for fdTMS and figure-8 coils. The results indicate that the fdTMS coils stimulate a smaller volume and are therefore more focal than their conventional counterparts for cortical targets at or closer to the surface than the *d*_½_ value, with all fdTMS coils being more focal than their conventional counterparts for target stimulation depths of less than about 1.1 cm in the brain. At a target stimulation depth of 0.5 cm, the fdTMS *d*_½_ ≥ 1.57 stimulates about half the volume of its figure-8 counterpart and about 30% less volume than the figure-8 coil with a target depth of 1.31 cm (Figure 10A, B). Furthermore, none of the three fdTMS coils would induce an E-field in the brain that is above twice the target threshold for targets 1.5 cm or closer to the surface (Figure 10C).

**Figure 10.**
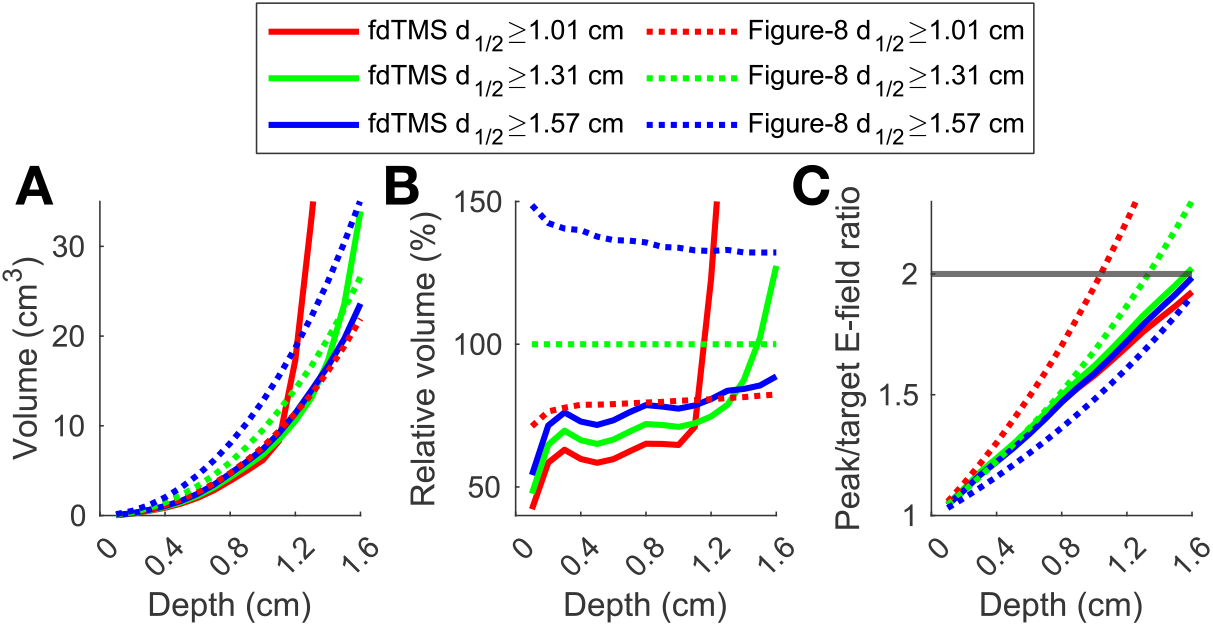
E-field distribution figures of merit versus target depth. (A) Total stimulated volume, (B) stimulated volume relative to the figure-8 *d*_½_ ≥ 1.31 cm coil, and (C) peak E-field in the brain relative to the E-field at the target at various depths in the brain. The depth corresponding to the intersection between the horizonal gray line and a curve is the value of the *d*_½_ figure of merit.

Figure 11 shows the coil energy relative to the figure-8 *d*_½_ ≥ 1.31 cm as a function of depth. While the energy of the fdTMS coils generally fall between the figure-8 coils with *d*_½_≥ 1.01 and 1.31 cm, the energy relative to the figure-8 *d*_½_ ≥ 1.31 cm coil decreases as target depth increases. At a target stimulation depth of 0.5 cm, the figure-8 *d*_½_ ≥ 1.01 coil and its fdTMS counterpart have similar energies. The other two fdTMS coils have slightly lower energies, but no lower than that of the figure-8 *d*_½_ ≥ 1.31 cm.

**Figure 11.**
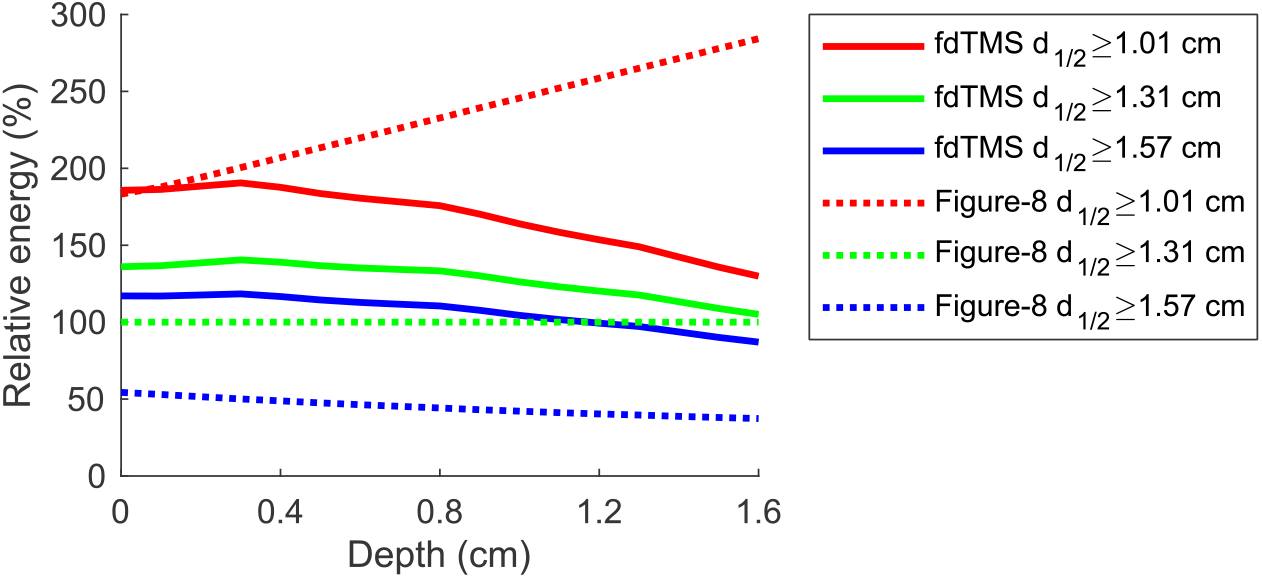
Energy of coils relative to the figure-8 *d*_½_ ≥ 1.31 cm coil.

TMS coils generate strong E-fields that are known to cause scalp sensations during stimulation. Figure 12 shows the scalp surface area stimulated above the target E-field and peak E-field relative to the target E-field. The fdTMS coils stimulate a significantly larger region of the scalp compared to the figure-8 coils, with the fdTMS *d*_½_ ≥ 1.01 cm coil stimulating approximately 5 times the scalp surface area compared to the figure-8 *d*_½_ ≥ 1.31 cm coil at a target depth of 0.5 cm, and the other two fdTMS coils stimulate approximately 3 times the scalp surface area (Figure 12A, B). On the other hand, the peak E-fields induced on the scalp are generally between those of the figure-8 *d*_½_ ≥ 1.01 cm and 1.31 cm coils (Figure 12C). While the peak E-field on the scalp is in line with conventional coils, the increased extent of scalp stimulation with fdTMS coils might result in more discomfort.

**Figure 12.**
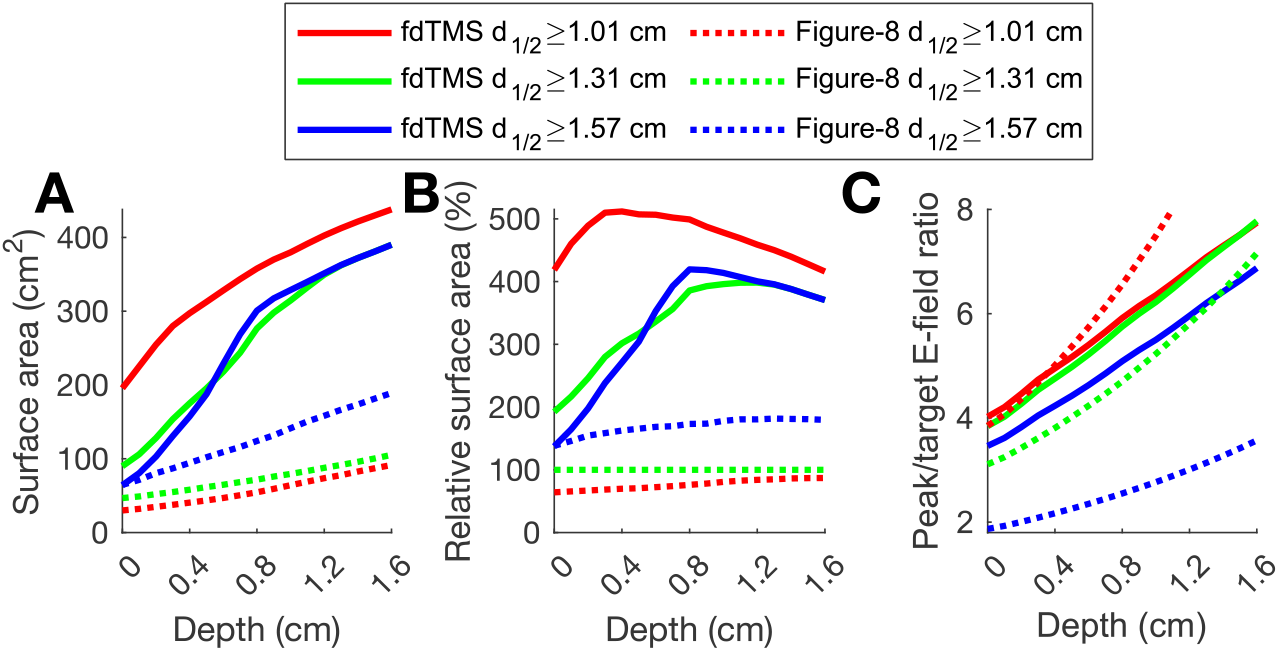
Characterization of scalp stimulation. (A) Total stimulated scalp surface area, (B) stimulated scalp surface area relative to the figure-8 *d*_½_ ≥ 1.31 cm coil, and (C) peak E-field on the scalp relative to the E-field at the target at various depths in the brain.

### Electric field measurements and validation of implemented coils

Ultimately, we implemented only the hybrid-layer coils with *d*_½_ ≥ 1.31 cm and *d*_½_ ≥ 1.57 cm, since our simulation evaluation indicated that the focality benefits of the *d*_½_ ≥ 1.01 cm design is restricted to very superficial layers. Figure 13 shows the final fdTMS coil implementations using the respective winding designs from Figure 7.

**Figure 13.**
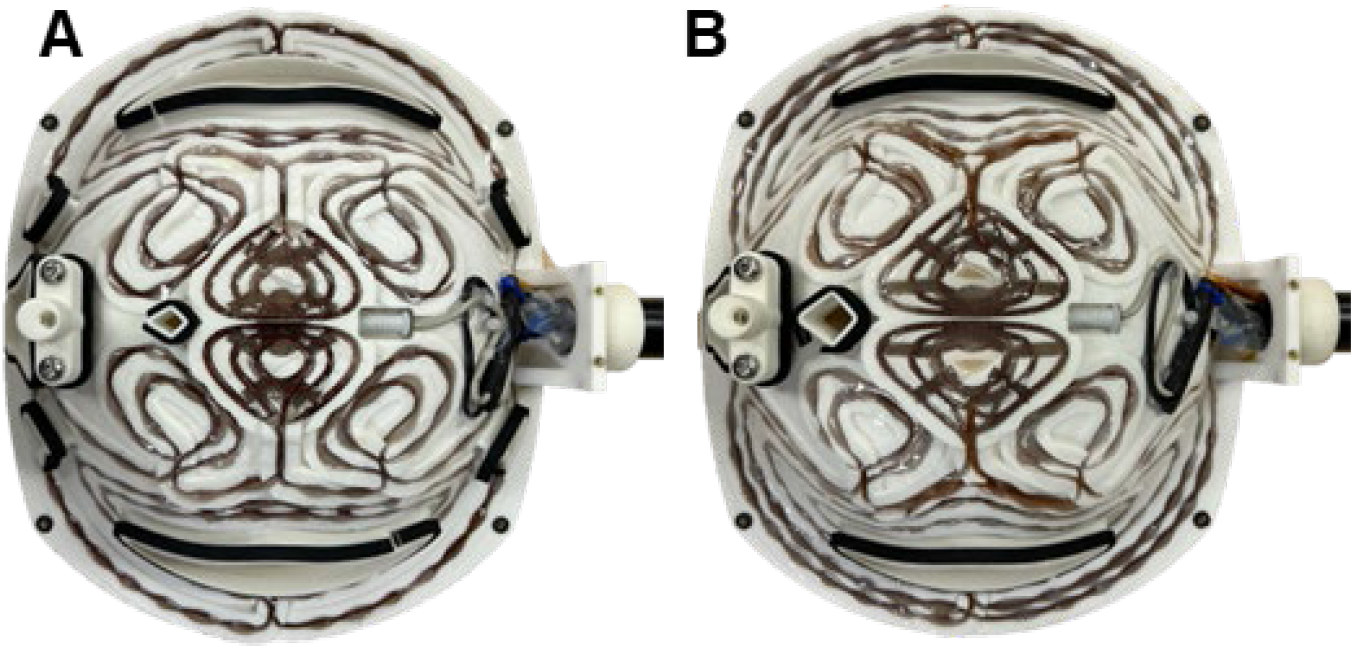
Hardware implementation of the hybrid-layer fdTMS coils with (A) *d*_½_ ≥ 1.31 cm and (B) *d*_½_ ≥ 1.57 cm and winding patterns shown in Figure 7. In addition to the windings, the coil formers house a neuronavigation tracker pedestal on the left and a temperature sensor (gray) and threaded handle base on the right.

The E-fields on a 7 cm and 6 cm hemispherical shell were measured for the fdTMS *d*_½_ ≥ 1.31 cm and *d*_½_ ≥ 1.57 cm coils as well as the commercial MagVenture Cool-B65 and B80 coils, which have the same respective depth figures of merit. The E-field measurement results are shown in Figure 14, indicating that the simulated and measured E-fields match well on both shells, validating the coil implementation. Furthermore, the contour enclosing the region with an E-field strength above 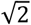 times its maximum is smaller for the fdTMS coils compared to their depth-matched figure-8 counterparts, demonstrating that the fdTMS coils are more focal.

**Figure 14.**
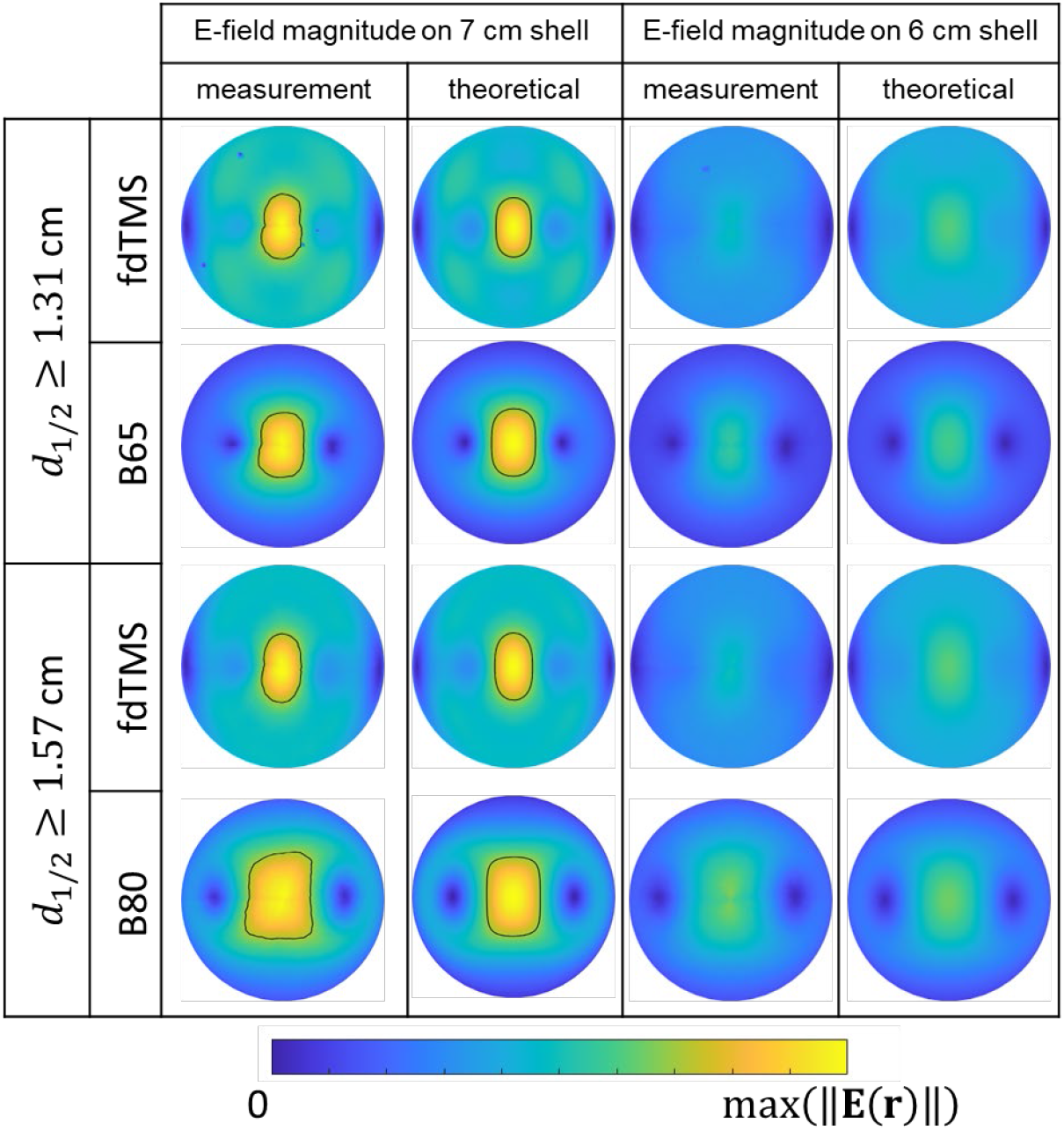
Coil E-field measurements on 7 and 6 cm hemispherical shells compared to theoretical E-field calculations. The black outline encloses the region with E-field strength above 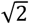 of the maximum E-field strength.

Figure 15 shows the measured E-field pulse waveforms generated by the experimental fdTMS coils and their conventional figure-8 counterparts. The pulse duration of both monophasic and biphasic pulses is comparable across the coils, due to the similar coil inductances. The fdTMS coils have more electrical damping, as evidenced by the lower ending amplitude of the biphasic pulses. Specifically, the energy loss is 77% and 76% for the *d*1/2 ≥ 1.31 cm and *d*1/2 ≥ 1.57 cm fdTMS coils, compared to 38% and 37% for the MagVenture B65 and B80, respectively. This is due to the larger number of windings in the fdTMS coils, resulting in longer current path through the winding wire, as well as the constraints on the wire diameter to accommodate the dense winding patterns in the center of the coil.

**Figure 15.**
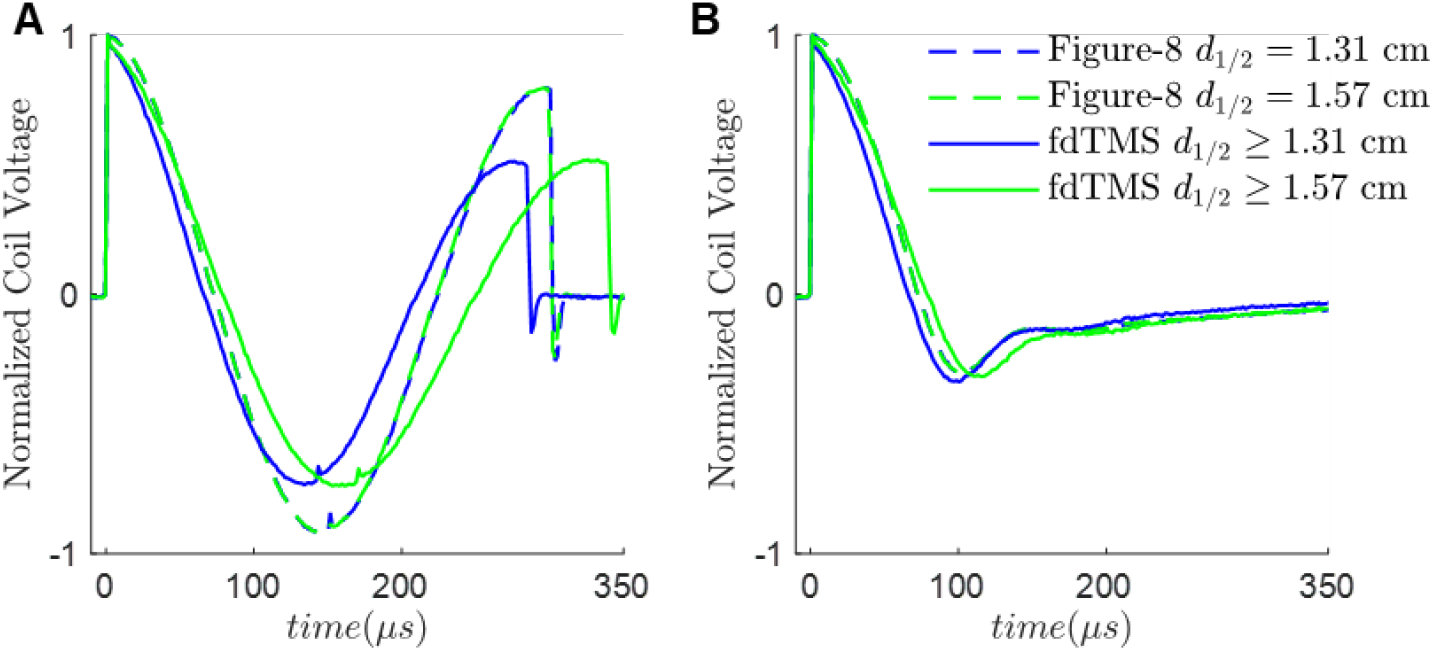
E-field pulse waveforms recorded from the experimental fdTMS coils with *d*_½_ ≥ 1.31 cm and *d*_½_ ≥ 1.57 and their figure-8 coil counterparts, MagVenture B65 and B80, respectively. Coils are driven with MagPro X100 with MagOption in standard pulse mode with (A) biphasic and

## DISCUSSION

We introduced a practical method for designing and fabricating fdTMS coils and validated the designs with experimental E-field measurements. The design framework includes computational optimization of the winding pattern, an energy-efficient curved “hat” coil surface enabling a wide range of placements, and a hybrid winding approach that improves energy efficiency by using multi-layer windings in some regions of the coil and a single layer in others. The E-field calculations, simulations, and measurements indicate that the fdTMS coils are 19–33% more focal than their depth-matched figure-8 coil counterparts at their respective target depth. The hybrid layer fdTMS coils achieve increased focality of 24–27% in a range of depths of 1.31–1.57 cm. For stimulation depth of 1.01 cm, the hybrid layer coil design achieved 17% increased focality and required 33% less energy than its the small figure-8 coil counterpart, indicating that fdTMS coils could also provide a more energy efficient alternative to small figure-8 coils. For a given coil, as the current is increased the depth of stimulation is also increased and focality decreases. The fdTMS coils outperformed figure-8 coils in terms of focality for all stimulation depths below the prescribed targeted stimulation depth. Correspondingly, the focality improvements are expected for a wide-range of coil driving current levels.

The design of focal TMS coils involves mediation with significant trade-offs between focality, depth, and energy. This is illustrated by the physics of TMS induced E-fields in a homogenous spherical head model (11). Equation 13 in reference (23) indicates that as the order of spherical harmonics on a spherical shell is increased, the decay will increase geometrically. In other words, sharp variations in the E-field on a spherical shell will result in rapidly decaying E-fields into the head. Furthermore, since higher order harmonics decay rapidly into the head, all of their associated energy is localized at shallow depths. This in turn results in large energy requirements to generate significant sharp E-fields deep into the brain.

The fdTMS coils produce a broader E-field distribution on the scalp than standard coils, indicating that a larger region of scalp tissue will be stimulated. One reason for this is that the fdTMS coils have large ‘biasing’ loops, which increase the E-field of large regions of the brain to near threshold. The E-fields associated with these ‘biasing’ loops are stronger on the surface of the scalp, thereby, resulting in broad regions of scalp exposed to strong electric fields. The impact on tolerability of the wider E-field spread on the scalp must be experimentally characterized. If tolerability is found to be a significant deterrent, additional constraint on the scalp E-field would have to be added to designs.

The resistive losses in the fdTMS coil are higher than in the conventional coils due to the presence of multiple loops and dense patterns near the coil center which collectively required a long stretch of winding wire and constrained the wire diameter to 2.05 mm. These levels of power loss would be acceptable for single-pulse applications like mapping, where monophasic pulses are commonly used. For repetitive TMS applications, where biphasic pulses are typically used and pulse energy recovery is important, the fdTMS coils will have to be redesigned to reduce the energy loss, which may result in reduction of their focality advantages.

## CONCLUSIONS

The fdTMS coil designs developed in this paper optimized focality within the constraints of brain stimulation by magnetic induction including limitations on the electric field depth, pulse energy, coil size, and winding density. These coils are more focal than their depth-matched figure-8 coil counterparts, and we believe these improvements represent the practical limits of E-field focality of TMS. Limitations of the fdTMS coils include the possibility of more discomfort associated with broader scalp stimulation as well as increased energy loss and associated heating. Overall, the presented design framework pushed the practical limits of TMS coil focality and identified potential advantages as well as challenges for future fdTMS applications.

## ACKNOWLEDGEMENT

Research reported in this publication was supported by the National Institute of Mental Health and the National Institute of Neurological Disorders and Stroke of the National Institutes of Health under Award Numbers RF1MH114268 and R01NS088674-S1. The content is solely the responsibility of the authors and does not necessarily represent the official views of the National Institutes of Health. The authors thank Dr. Moritz Dannhauer for guidance on E-field modeling and Eleanor Wood for help with coil construction.

